# Predicting kinship dynamics during pre- and post-reproductive life stages

**DOI:** 10.1101/2025.04.11.648073

**Authors:** Peng He, Michael N. Weiss, Samuel Ellis, Daniel W. Franks, Michael A. Cant, Darren P. Croft, Rufus A. Johnstone

**Author notes:** Correspondence (PH), (RAJ).

## Abstract

Individuals’ relatedness to their groups often changes with age, potentially favouring age-linked trends in social behaviours, and these kinship dynamics have been invoked to explain the evolution of life history traits such as early reproductive cessation and post-reproductive helping. While models showed that simple demographic parameters (e.g., the rates of male and female philopatry vs dispersal, and of local vs non-local mating) suffice to predict the patterns of kinship dynamics breeders experience, they make the unrealistic assumptions that survival and fecundity are age-independent, thus fail to capture the full complexity of kinship dynamics. In particular, they cannot consistently predict how individuals’ relatedness to their groups should change during their pre- and post-reproductive phases. Here, we extend existing models of kinship dynamics to allow any age-linked changes in mortality and fecundity of both sexes. We describe the new model, and demonstrate its improved predictive power, by comparing the observed kinship dynamics of the killer whales to the predictions of the previously available models (assuming constant mortality and fecundity) and of our new model (considering any observed age-specific survival and fecundity). We show the new model better predicts the data, successfully capturing the observed ‘three-stage’ female kinship dynamics: an individual’s relatedness to her group decreases initially while juvenile, increases across reproductive lifespan, and decreases again during post-reproductive life. These predictions demonstrate the power of our model in generating new insights into the theory of social life history evolution (e.g., explaining why post-reproductive lifespan evolved yet are constrained).

## 1 INTRODUCTION

There is clear empirical evidence that in some species, the average relatedness of an individual to others in its social group changes systematically across its lifespan, often in ways that differ for females and males (Croft et al., 2021). These sex-specific kinship dynamics have been invoked to explain age-related trends in the expression of social behaviours such as helping and harming (Johnstone and Cant, 2010, Ellis et al., 2022). For example, in contrast to males, female killer whales exhibit early reproductive cessation followed by a significantly extended post-reproductive lifespan during which they assist kin (Croft et al., 2017, Ellis et al., 2024). This phenomenon of post-reproductive helping, which is rare among mammals (Ellis et al., 2018, 2024), has been linked to the unusual kinship dynamics these females experience — their average relatedness to others in their social groups typically increases with age across the reproductively active portion of the lifespan (Croft et al., 2017, Ellis et al., 2022), a pattern which is thought to be favourable to the evolution of menopause and late-life helping (Johnstone and Cant, 2010, Croft et al., 2017, Johnstone and Cant, 2019, Ellis et al., 2022).

To understand how kinship dynamics emerge in social groups, as well as the inclusive fitness effects they can have on social behaviours expressed at different life history stages, models have been constructed to investigate how demographic parameters, such as the rates of female and male philopatry versus dispersal and the rate of local versus non-local mating (Johnstone and Cant, 2008, 2010, Ellis et al., 2022), influence sex- and age-specific relatedness. While these models offer considerable insight into the emergence of kinship dynamics in animal societies characterized by diverse patterns of dispersal and local mating, their predictions typically rest on the simplistic assumption that fecundity and mortality of the sexes are age-independent (i.e., constant across lifespan). In natural populations, however, these demographic parameters are typically age-linked (Pianka and Parker, 1975, Keyfitz and Caswell, 2005, Lee et al., 2016) and may be expected to shape kinship dynamics individuals experience — such as increased mortality by ivory poaching targeting at (adult) females in specific age classes amplifies the age-linked changes of a female African elephant’s relatedness to its maternal family kin (Croll and Caswell, 2025).

Here, we extend existing models of kinship dynamics (Johnstone and Cant, 2010, Ellis et al., 2022, He et al., 2023) to explicitly incorporate the age dependencies of survival and fecundity for both sexes. We first describe the new model, and then demonstrate its advantages over previous analyses in predicting key qualitative features of kinship dynamics, using as an illustrative example of the patterns of kinship dynamics observed in southern resident killer whales. In particular, we show that by contrast with its more simplistic predecessors, the extended model successfully predicts the kinship dynamics experienced by females not only during an individual’s reproductive span (where they are increasingly related to their groups), but also during pre- and post-reproductive life stages (where their relatedness to their groups declines). By being able to capture the full complexity of kinship dynamics, our model offers new insights into social life history evolution.

## 2 THE MODEL

### 2.1 Demographic Assumptions

Following Johnstone and Cant (2010), Ellis et al. (2022), we consider an infinite, diploid, bisexual population divided into discrete social groups (i.e., the infinite island model, Wright, 1931). Time proceeds in discrete steps — at each timestep there are *N*_*α*_ females and *N*_*β*_ males in each group (i.e., at each timestep each group has *N*_*α*_ + *N*_*β*_ breeding vacancies). In notations below, females and males are indicated by *α* and *β*, respectively. The age class of an individual with sex *ϕ* ∈ {*α, β*} in a group is denoted by *K*_*ϕ*_ (the sex is implicitly indicated by *ϕ* here and *ψ* later), and where ℕ^+^ is the set of positive natural number, *K*_*ϕ*_ = 1 and *K*_*ϕ*_ = *C*_*ϕ*_ are the lowest and highest age class of the sex *ϕ*, respectively, and 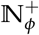 is the set of age class for the sex *ϕ*.

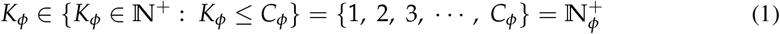

As time elapses, the female and male age class compositions in a group can change following the demographic processes assumed. We model such transitions using a finite discrete-time Markov chain, where the state of a group at each timestep is uniquely identified by a non-decreasing age class sequence of females followed by that of males. With given *N*_*α*_, *N*_*β*_, *C*_*α*_, *C*_*β*_, we denote the number of states in which a group can be found at any timestep as *N*_*s*_ — i.e., the size of group state space denoted by the finite set Ω ≠∅. Group state transition probabilities depend on the sex- and age-specific survival probabilities of group members. We denote the probability that an individual *i* with sex *ϕ* in a group (in any state) survives to the next age class (i.e., 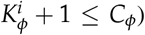 as 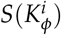, where 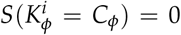 indicates the individual cannot survive beyond its life expectancy. Note that individual identity such as *i* in a group is introduced for clarity, but survival rates, as well as fecundity below, are sex- and age-class-specific thus independent of *i*.

We assume female population demographic dominance, where males can always fertilize all the females in the population and the number of offspring produced depends only on the age-specific fecundity of females (de Vries and Caswell, 2019). The age-specific fecundity of a male is the amount of paternity he enjoys over the offspring reproduced in the population — by competing with other males. At a given timestep in a group in state *s*, the number of offspring a female *i* with age class 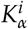 reproduces relative to the average of all the female age classes, or the number of offspring a male *i* with age class 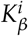 sires relative to the average of all the male age classes, is denoted by 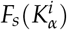 or 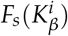, respectively (i.e., the sex and age-class-specific fecundity). We assume that the absolute values of 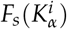 and 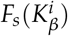 are large enough so that the breeding vacancies in a group are always fully occupied, and that at each timestep a proportion ζ of the offspring produced by each female in a group are females while the rest are males (this primary sex ratio does not influence predicted kinship dynamics but are introduced for clarity), and that of all the offspring produced in a group, a proportion *ρ* are sired by local males while 1 − *ρ* by non-local males. An individual *i* sexed *ϕ* in age class 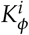 survives to its next age class with the probability 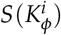 described earlier, while an offspring sexed *ϕ* disperses to a random non-natal group with the probability *d*_*ϕ*_. After dispersal, offspring in each group (both those born locally that did not disperse, and those immigrants from elsewhere in the population) compete in a fair lottery for the breeding vacancies left by deaths of others of their own sex in that group, and those failed to obtain any breeding vacancy die, and the cycle then repeats.

### 2.2 Group Steady-state Distribution

With given sex- and age-class-specific survival probabilities and the group state space Ω, we define an *N*_*s*_ × *N*_*s*_ Markov transition matrix ***T*** — the entry 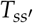 of which is the probability that the group transits from *s*^′^ to *s* in a timestep (calculated with the survival probabilities of the group members), and

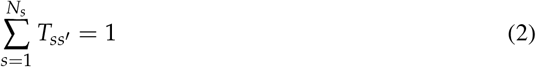

(i.e., ***T*** is row-stochastic such that each of its rows sums up to 1). With ***T***, we can derive the stationary probability *u*_*s*_ that a group is found in state *s* at any given timestep after the population reaches its demographic equilibrium, by solving for the dominant left eigenvector ***u***_*s*_ of ***T*** (i.e., the row vector of length *N*_*s*_ associated with *λ* = 1 known as the dominant left eigenvalue; Philippe et al., 1992) with

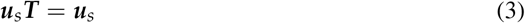

Note that with larger group size and/or the numbers of age classes for the sexes, the size of ***T*** may demand more computational capacity than practically available. For such cases, we also describe (in the APPENDIX) an analytical solution for (3), where ***u***_*s*_ can be alternatively derived from the stable age class distributions of the sexes (under the sex- and age-class-specific survival probabilities of interest).

### 2.3 Summary Statistics and Probabilities

With the demographic processes assumed, at each timestep the expected number of offspring produced in a group is

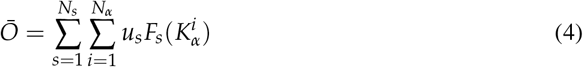

while the average number of immigrants sexed *ϕ* ∈ {*α, β*} to a group at each timestep is

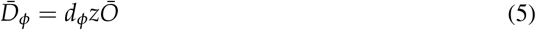

where *z* =ζ when *ϕ* = *α* (females), while *z* = 1 −ζ when *ϕ* = *β* (males). Then, the probability that an individual sexed *ϕ* ∈ {*α, β*} is born to a local female *i* with age class 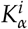 in a group with current state *s* is

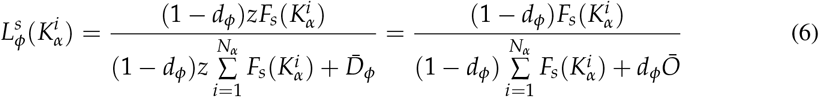

while the probability that it is native to the group (i.e., born to any female in the group) is

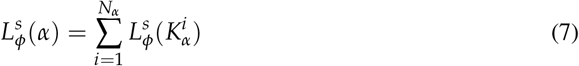

For males, the expected number of offspring they sire in a group at each timestep is

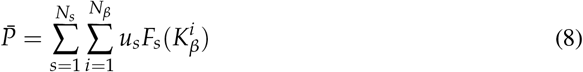

and of these offspring, an expected amount

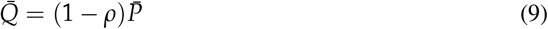

are born elsewhere in the population, and thus the probability that an offspring born in a group with state *s* is sired by a local male *i* with age class 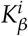 is

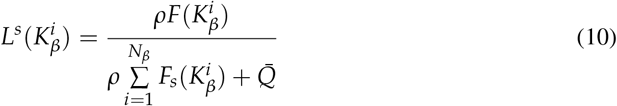

Under the demographic assumptions, at any given timestep, the probability that an individual sexed *ϕ* is found in the age class *K*_*ϕ*_ in the population is

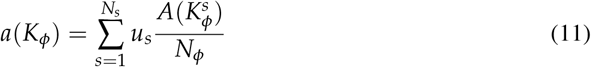

where 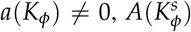 is the number of the age class *K*_*ϕ*_ in the group in state *s*, and *N*_*ϕ*_ is the number of individuals of the sex *ϕ* in each group satisfying 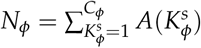.

### 2.4 Probabilities of Inheritance

In a group with current state *s*, the probability that a selectively neutral homologous gene will be passed on from a female *i* with age class 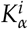, or a male *i* in age class 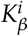, to an individual sexed *ϕ* ∈ {*α, β*} native to the group at the next timestep is

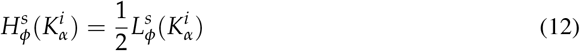

or

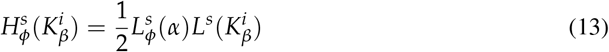

respectively.

### 2.5 Kinship Dynamics

We model kinship dynamics based on the Markov process described above. Relatedness between individuals is defined as the probability of *identity by descent* (IBD) of two above-mentioned gene copies, each of which is randomly sampled from each of the two individuals (Johnstone and Cant, 2008, Ellis et al., 2022). In calculations, we use a symmetric matrix to capture the pairwise relatedness in a group at each timestep. Initially, we assume that individuals in a group (in any state) are not related to others — as captured by the (*N*_*α*_ + *N*_β_) × (*N*_*α*_ + *N*_*β*_) identity matrix ***R*** (note the memoryless property of this Markovian-based approach implies that the predicted kinship dynamics is independent of the initial relatedness assumed). As time elapses, inter-individual relatedness in a group builds up and eventually converges (such equilibrium is guaranteed by the underlying Markov process and is independent of the initial values, Levin and Peres, 2017), and the expected relatedness between individuals in a group in the current state *s* can be expressed as

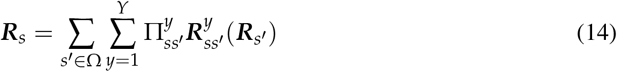

In (14), *Y* is the number of unique sequences of survival and death events as the ‘fates’ of the females and males in the group that was in state *s*^′^ at the last timestep. 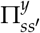 is the probability that the group state transition to *s* was realized by a particular fate sequence *y*, given the sex- and age-class-specific survival probabilities of the individuals (here 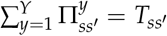 and 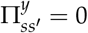 indicates the transition from *s*^′^ to *s* could not be realized by *y*). 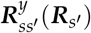 is an (*N*_*α*_ + *N*_*β*_) × (*N*_*α*_ + *N*_*β*_) matrix capturing the expected relatedness between individuals following the group transition from *s*^′^ to *s* via *y*, where 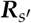 captures inter-individual relatedness when the group was in state *s*^′^.

When deriving ***R***_*s*_ following (14), we first classify each pair of individuals by their sexes and whether they are newly-established breeders or not after the transition to *s*. Specifically, for a given fate sequence *y* realizing the transition, we use 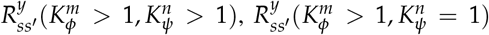, or 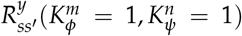 to denote the expected relatedness between individuals identified as *m* (sexed *ϕ*) and *n* (sexed *ψ*), where newly-established breeders are indicated by 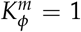 or 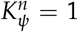 while survivors by 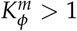 or 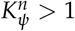. Then, we calculate the expected relatedness between *m* and *n* at the current timestep using (12), (13), and/or 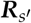 from the previous timestep, with

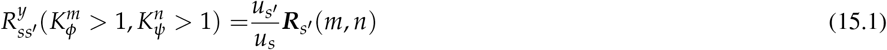

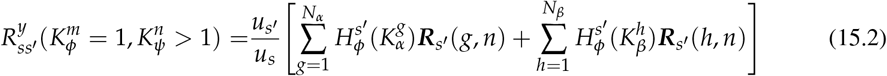

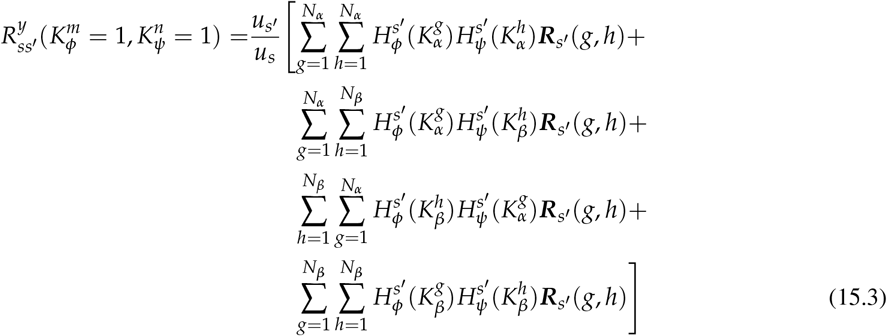

With initial 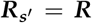, we can iterate (14) with (15.1), (15.2), and (15.3), until the most dramatic increment of pairwise relatedness (e.g., 0 *< ϵ* ≪ 1) to ***R***_*s*_ becomes trivial. Then, for an individual sexed *ϕ* ∈ {*α, β*} in age class *K*_*ϕ*_, its average relatedness to others with sex *ψ* ∈ {*α, β*} is

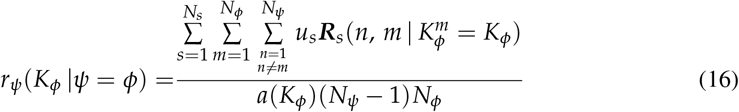

if others are with the same sex (*ϕ* = *ψ*), or

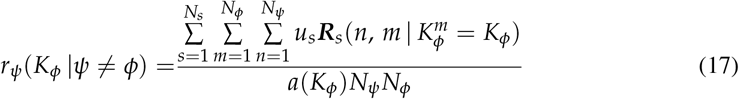

if others are with the opposite sex (*ϕ* ≠ *ψ*), where 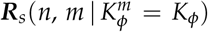 captures the relatedness between the focal individual *m* sexed *ϕ* in age class *K*_*ϕ*_ (if any) and its group mate *n* sexed *ψ* in a group with state *s*. Lastly, the average relatedness of an individual sexed *ϕ* ∈ {*α, β*} with age class *K*_*ϕ*_ to its group is calculated as

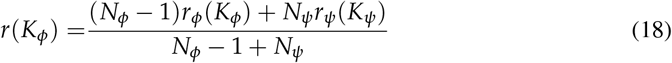

where *ϕ* ≠ *ψ*.

### 2.6 A Note for a Markovian-based Alternative Approach

In the analytical approach described above, group state space (*N*_*s*_) expands rapidly with group size and/or the numbers of age classes for the sexes, and this may pose a computational challenge for relatedness calculations encapsulated in (14), especially for species with large group size and/or long lifespan (e.g., killer whales, see example below). In such scenarios, while we can define less group states so that relatedness calculations following (14) is computationally not constrained (e.g., by aggregating each consecutive multi-year age interval into an age class instead of using natural ages), doing so would incorporate less explicit demographic info into the model (e.g., the average survival and fecundity at each multi-year age interval instead of such demographic parameters explicitly observed at each age), and this does not allow us to predict the fine-scale patterns of kinship dynamics (e.g., how average relatedness to a group changes with an individual’s natural age).

To address such computational challenge while still being able to make explicit use of demographic info for predictions, we also describe (in the APPENDIX) an alternative Markovian-based ‘hybrid’ version of our analytical model captured in (14). In this hybrid approach, the patterns of kinship dynamics are derived from the analytically *expected* inter-individual relatedness under *simulated* changes of the age (class) compositions of a focal group over time, with the given sex- and age- (class-)specific survival probabilities of interest. As it considers only the realized/simulated age (class) composition of the focal group at each timestep (i.e., a simulated instead of an analytically defined Markov process — the latter of which considers all the possible group state transitions at each timestep), its relatedness calculations are computationally far less demanding than those would be while following (14) analytically. With this hybrid alternative, we are thus able to not only consider realistic group sizes (which modulate kinship dynamics, He et al., 2023), but also incorporate demographic parameters as explicitly as they are empirically available for predictions — as shown in the killer whale example below, the hybrid approach allows incorporation of the *age-explicit* survival and fecundity for predictions, instead of such demographic info aggregated at coarser age intervals (e.g., those consecutive five-year age windows as age classes for these long-lived group-living creatures — see details in the APPENDIX) we would otherwise need to define to reduce group state space to such an extent that with our given computational capacity calculations following (14) for this species is feasible (we show in the APPENDIX the predictions by our analytical model at such coarser five-year age intervals as age classes for the whales).

## 3 RESULTS

In this section, we compare the observed kinship dynamics experienced by southern resident killer whales of both sexes to those predicted by: (1) a model that assumes constant mortality and fecundity rates, as in previous studies (Johnstone and Cant, 2010, Ellis et al., 2022, He et al., 2023), and (2) an ‘improved’ model that incorporates sex- and age-specific changes in mortality and fecundity, analysed using the ‘hybrid’ Markovian-based approach described above. In the rest of this section, we refer to these as the ‘old’ and ‘new’ models, respectively.

### 3.1 Estimating model parameters and generating predictions

Fig. 1 shows, in panels A and B, the sex- and age-specific survival rates and relative fecundities assumed in the new model (solid dots joined by solid lines, Fig. 1-a, 1-b), which are derived from the empirically estimated demographic schedules for females and males in this population (Ford et al., 2018, Nielsen et al., 2021), and the corresponding, age-invariant survival rates assumed in the old model (open circles joined by dashed lines, Fig. 1-a). These latter, constant survival rates were chosen to ensure that both models feature the same, consistent mean lifespans for males and females (see APPENDIX for further details). Note that in the old model, the fecundities of both sexes are also assumed to remain constant across the lifespan; but as the predicted patterns of kinship dynamics depend only on relative fecundity at different ages, it is not necessary to estimate absolute fecundity, and no value is therefore shown for the old model in Fig. 1-b. The lower panels of Fig. 1 show the stable age distributions that result from the assumptions of the two models, with mean lifespans for both sexes indicated by vertical dashed lines.

**Fig. 1.**
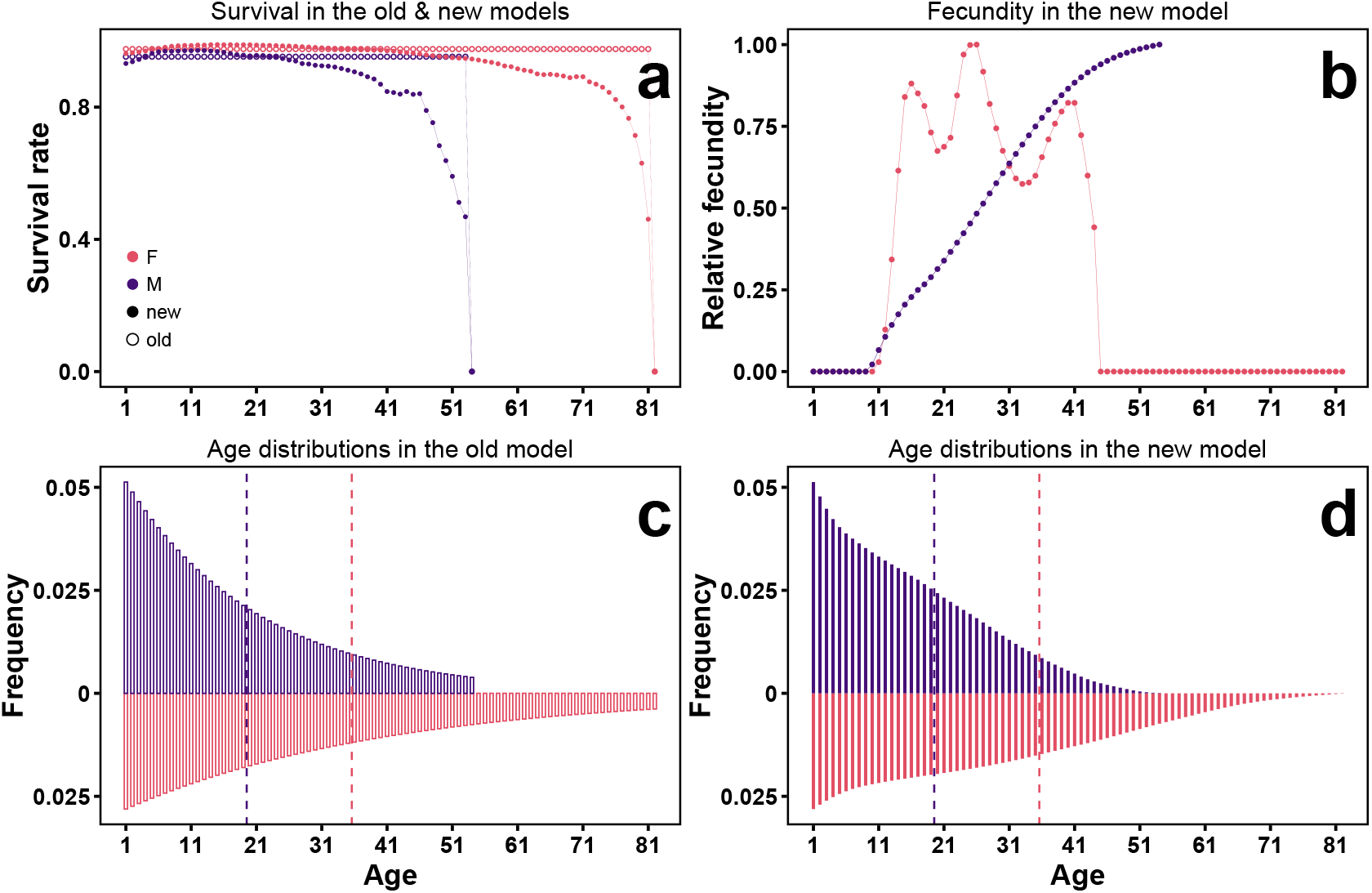
The (sex- and) age-specific (a) survival rates and (b) relative fecundity (scaled by the ‘most’ fecund individual in each sex) of the southern resident killer whales (‘F’ and red: female; ‘M’ and purple: male) as demographic inputs for the current Markovian-based models of kinship dynamics (solid dots joined by solid lines), in contrast to the counterpart age-independent survival rates for the sexes as inputs for the previous models (open circles joined by dotted lines). The constant survival rate for a sex is derived as that that gives the same mean lifespan as the age-specific survival rates would do — as marked by the dashed vertical lines for the sexes in (c) and (d) showing the stable age distributions these survival rates would give for the sexes (unfilled bars in c by constant survival probabilities; filled bars in d by the age-specific survival probabilities; see in the APPENDIX for the derivations of these demographic inputs for both models). Note that it is not necessary to estimate the precise constant fecundities assumed for the sexes for the previous models, as their predictions are independent of these constants (their predictions depend on the relative fecundity across ages).

Since mating in the southern resident killer whale population typically occurs between distinct groups and both sexes remain within their matrilineal group (Bigg et al., 1990, Ellis et al., 2022), we assume in both models that *ρ* = 0.02 and *d*_*α*_ = *d*_*β*_ = 0 (as Ellis et al., 2022).

The only remaining parameters in either model are the numbers of females and males per group (i.e., *N*_*α*_ and *N*_*β*_). As killer whale matrilineal group sizes exhibit considerably variations, there is no consistent group size as input for either model. Thus, for each model separately, we calculated the predicted mean relatedness of males and females to their social group across the lifespan, for all possible combinations of *N*_*α*_ and *N*_*β*_ taking integers from 1 to 7 (i.e., the smallest or largest group modelled has 2 or 14 individuals, respectively, where sex ratio ranges from 1:7 to 7:1; Fig. A1). Results shown below are those for the ‘best-fit’ numbers of females and males for the model in question, which minimise the sum of squared deviations of predicted relatedness values across both sexes and all age classes from those observed in individual whales at each age. While the ‘best-fit’ number of males (i.e., 3) is the same for both models, the ‘best-fit’ number of females is higher for the new than for the old model (i.e., 6 rather than 5; Fig. A1). This difference reflects the fact that the new model incorporates age-specific variation in female fecundity, in particular due to the occurrence of pre- and post-reproductive individuals, which lowers the effective number of breeding females within a group.

Predicted mean relatedness values were derived analytically for the old model, following the approach of He et al. (2023). For the new model, we used the hybrid approach described in the previous section, tracking age-specific mean relatedness values within each of 50 simulated chains over the final 10^4^ of a total 10^6^ time-steps (i.e., with a burn-in period of 9.9 × 10^5^ time-steps in each chain) and taking the means of these age-specific values across the chains as the model predictions (see DATA & CODE AVAILABILITY and APPENDIX for further details).

### 3.2 Model predictions

Fig. 2 shows the observed mean relatedness of male (upper panel) and female (lower panel) resident killer whales to their group, as a function of age, alongside the predictions of the old model (in blue) and the new model (in red, where the standard error associated with each predicted mean value was, for all age-classes in both sexes, less than 4 × 10^−4^). The two-sample Kolmogorov-Smirnov (KS) test suggests that, with the respective ‘best-fit’ numbers of females and males for each model, the predicted sex- and age-specific relatedness values by the new model fit significantly better to those empirically observed than the previously available model does (i.e., the cumulative distribution of the squared deviations of the ‘new’ predictions from observations is significantly different from the cumulative distribution of the squared deviations of the ‘old’ predictions from observations; KS test statistic *D*=0.076, *p<*0.001; the sum of squared deviations for the ‘new’ and ‘old’ model: 11.5 and 11.9, respectively; see Fig. A2 in the APPENDIX for the distributions).

**Fig. 2.**
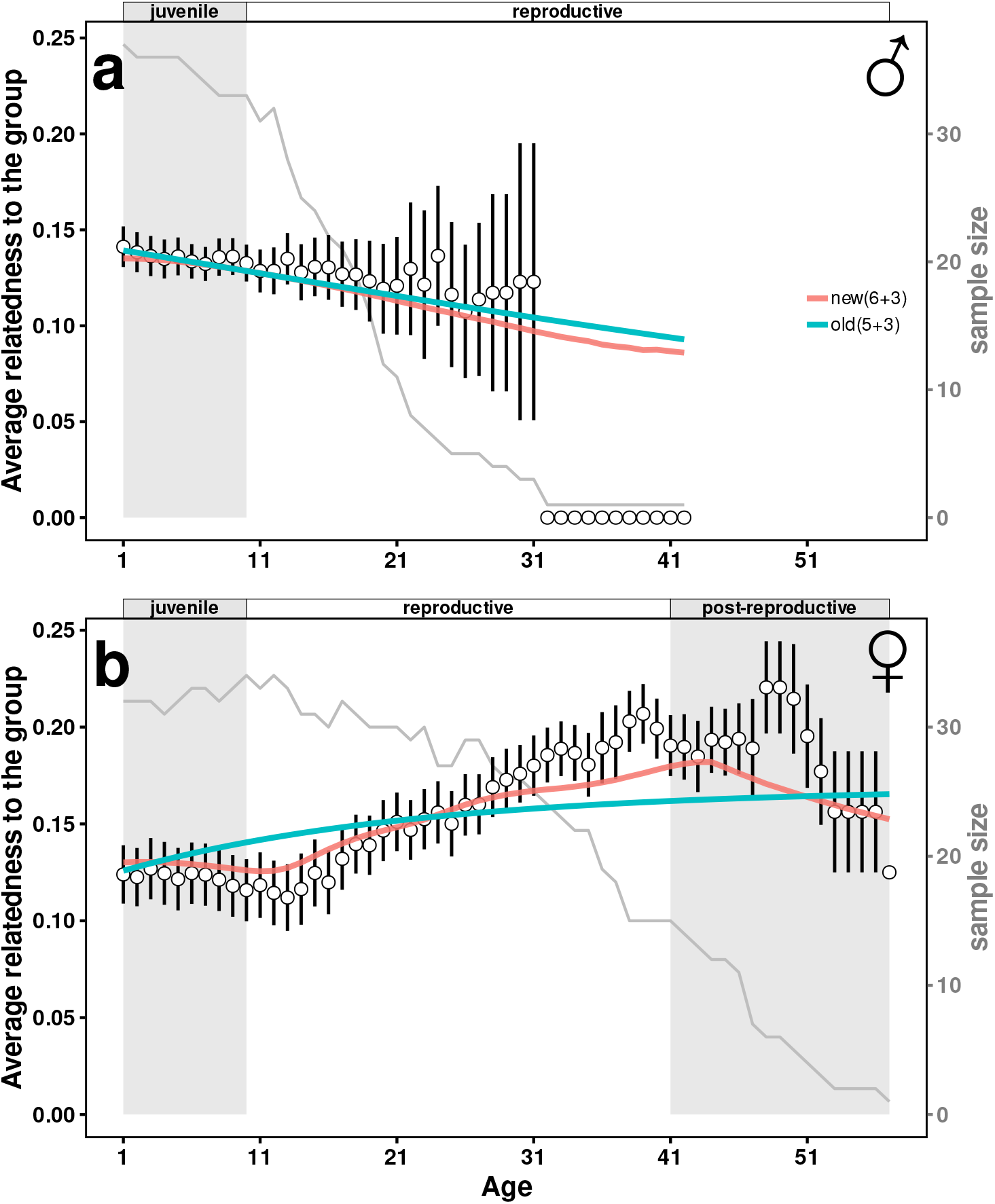
The empirically observed patterns of kinship dynamics experenced by the southern resident male (a) and female (b) killer whales (i.e., circles with bars indicating the standard error of the mean age-specific average relatedness of individuals to their matrilineal groups), along with their ‘best-fit’ predictions by: (1) the old model (blue) that was previously available, which takes the age-independent survival and fecundity for both sexes, and (2) the current new model (red), which allows age-specific survival and fecundity for both sexes (note the standard error bars indicating the accuracy the predictions by the new model are too small to be visible). Here, a group of 5 females and 3 males in the old model, or a group of 6 females and 3 males in the new model, gives rise to such a set of predictions that best explains the observed sex- and age-explicit average relatedness of individuals to their groups (i.e., individual-based observations). The grey lines show the age-specific sample sizes for the sexes in the relatedness data (from which the observed means and standard errors are derived), while the grey shaded age intervals mark the non-reproductive life history stages of the sexes.

As Fig. 2-a reveals, incorporating age-linked changes in mortality and fecundity has minimal impact on predicted kinship dynamics for male killer whales — both models generated 31 (out of 42) age-specific predictions falling within the standard error of the observed means. However, for female killer whales, the incorporation of the observed age-explicit changes of survival and fecundity in the new model predicts different fine-scale patterns of kinship dynamics than those by the previously available old model (Fig. 2-b) — across the 57 age-specific observations, the new and old models generated 44 and 25 predictions, respectively, that fall within the standard error of the observed means (Fig. 2-b). In particular, while the previously available model predicts that a female’s relatedness to her group should increase monotonically (though asymptotically) with age, the new model predicts a marked decline in average relatedness with age for post-reproductive females. Likewise, the fine-scale patterns of kinship dynamics predicted by the new model for female juveniles are more similar to those empirically observed than the predictions by the old model are: the new model realistically captures the decreasing trend of a pre-reproductive female’s relatedness to her group, while the previously available model would instead predict an opposite trend (Fig. 2-b).

## 4 DISCUSSION

Earlier models of kinship dynamics have generated deep insights into social life history evolution (Johnstone and Cant, 2010, Ellis et al., 2022, He et al., 2023), but none is able to capture the full complexity of kinship dynamics that may arise across the cycle of life under various demographic processes, thus limiting our general understanding of the evolution of sex- and age-linked trends in social behaviours. We address this gap by proposing a more general model that can incorporate the effects of sex- and age-specific survival and fecundity into predictions (while keeping considering the effects of dispersal and mating). With long-term empirical data, we demonstrated the improved predictive power of the model over its predecessors, and show that it successfully captures the diverse patterns of kinship dynamics female killer whales experience across life — not only during reproductive but also during pre- and post-reproductive stages. By being able to capture the full patterns of kinship dynamics individuals of both sexes experience, the model thus helps to stimulate new hypotheses and insights into social life history evolution.

As expected, we show that individuals’ average relatedness to their groups is sensitive to age-linked changes in survival and fecundity, thanks to the roles of these demographic schedules in shaping the gene genealogy within populations (Goodman et al., 1974, Pullum, 1982, Hudson, 1990, Donnelly and Tavare, 1995, Hammel, 2005, Metcalf and Pavard, 2007, Johnstone and Cant, 2010, Neher and Hallatschek, 2013, Wakeley et al., 2016). In our killer whale example, the predicted and observed ‘three-stage’ female kinship dynamics suggest that, in social groups where generations overlap and individuals are closely related to one another (i.e., no immigration of stranger and emigration of kin), the imbalances between the births and deaths of groupmates (kin) fundamentally drives the dynamics of an individual’s genetic ties to its group. For example, all else being equal, a focal female killer whale’s average relatedness to her group will remain unchanged if the death of her first-order relative (e.g., mother) is immediately followed by the birth of a relative of the same order (e.g., son or daughter — both of which are philopatric in this species); however, if the female is sexually immature or menopausal and the death is instead followed by the birth of a second-order relative (e.g., nieces/nephews), such imbalances are created for her and she becomes less related to her group. The age-linked changes in survival and fecundity play a fundamental role in creating such imbalances as individuals age — by predisposing the losses and gains of different types of relatives and/or their frequencies — if any, along with the systematic changes in the numbers of non-relatives/immigrants in a local group prescribed by both the survival schedules and the patterns of dispersal of the sexes (our model can capture all these effects). Interestingly, we note that in such kin groups, similarity in survival and fecundity schedules can create qualitatively similar patterns of kinship dynamics. For example, female killer whales and female African elephants both live in (matrilineal) kin groups and share similar survival and fecundity schedules — both are characterized by the typical type-I survival curves and their reproductive rates typically peak at middle ages (Lee et al., 2016, Nielsen et al., 2021); by analysing the changes of a focal female African elephant’s kin networks, it has been shown that her average relatedness to her kin group decreases before sexual maturity, increases as soon as she starts to reproduce, and decreases again when entering into elderhood (Croll and Caswell, 2025), as the ‘three-stage’ pattern of kinship dynamics we revealed.

While our new model enables us to predict the full patterns of kinship dynamics across whole lifespans of individuals in social systems characterized by diverse patterns of dispersal and local mating, we also note here the empirical challenges that may arise when generating and/or testing predictions. For example, in species with a typical type-I survival schedule (e.g., social mammals like the killer whales and our own species), it is common to see (in both demographic and relatedness data) that sample sizes for the older are increasingly smaller than those for the younger, and this can lead to greater discrepancies between predictions and observations at later than earlier life history stages (which thus highlights the value of long-term and systematic observational data in addressing such challenges). Despite such empirical challenges, we suggest that the predictions by our new model are generally more plausible than those by its predecessors, and while previous models may perform qualitatively equally well as our new model (e.g., when predicting the patterns of male killer whale kinship dynamics), the new model is generally more applicable — it allows us to incorporate as explicit survival and fecundity data as empirically available to account for the effects of the age-dependencies of these demographic parameters on the emergent patterns of kinship dynamics.

Given that kinship dynamics can impact the inclusive fitness outcomes of social behaviours expressed at different life history stages (Johnstone and Cant, 2010, Ellis et al., 2022, He et al., 2023), predictions by our new model thus can be invoked to stimulate new hypotheses and insights into sex- and age-linked trends in social behaviours. For example, under the theory of kin selection, the predicted (and observed) dramatic declines of post-reproductive female killer whales’ average relatedness to their groups may expect increasingly weaker selective pressure for these females to be ‘helpful’ for their groups as they age. Such a declining trend in kinship dynamics may also help to explain why there is eventually an ‘end’ of post-reproductive lifespans of these females: our supplementary analyses (Fig. A3) show that a (hypothetical) post-reproductive female dies at age 48 (i.e., ca. 7 yeas after entering menopause) is predicted to be as strongly related to her group as she was alive at the age of 35 — which is the (weighted) average age at death for females given the assumed stable age distributions. Likewise, the predicted (and observed) declines of pre-reproductive female killer whales’ average relatedness to their groups may instead expect increasingly stronger selective pressure for these females to be ‘harmful’ as they approach sexual maturity.

## DATA & CODE AVAILABILITY

Details on demographic and relatedness data analyses are enclosed in the appendix. All the code and data used are openly available at https://github.com/ecopeng/Model_Kinship_Dynamics.

## AUTHORS’ CONTRIBUTIONS

PH and RAJ led the conceptualization, modelling, and writing for this study with supports and inputs from other co-authors. MNW, SE and DPC led the curation, analysis, and interpretation of the killer whale relatedness data. DWF, MAC, DPC and RAJ contributed to acquiring funding for this study. All authors gave final approval for the publication of this study.

## ACKNOWLEDGEMENTS

We thank the Natural Environmental Research Council for granting us the funding (No. NE/S010327/1) to conduct this study.

