## APPENDIX for "Predicting kinship dynamics during pre- and post-reproductive life stages"

### CONTENTS


1 Age-dependent survival and fecundity 2

2 Constant survival and fecundity 2

3 The hybrid version of the current model 3

4 Calculating relatedness of killer whales 6

5 Supplementary figures 7

### 1 Age-dependent survival and fecundity

The sex- and age-dependent survival and fecundity of the southern resident killer whales are derived from the survial and fecundity schedules empirically estimated by two studies. From Nielsen et al. (2021), we get (i) the standardized survivorship curves for both sexes (the original Fig. 2-a), and (ii) the females' fecundity schedule (the original Fig. 3-a), while from Ford et al. (2018), we get (iii) the males' fecundity schedule (the original Fig. 2). We extracted these survival and fecundity schedules using the web-based tool *WebPlotDigitizer* (Rohatgi, 2022). The age-specific survival and fecundity are derived as their estimates by the local polynomial regression model fitted to the raw data extracted (see code availability for details). The mean survival and fecundity in each (5-year) age class for each sex are derived (as the inputs for the new analytical model of kinship dynamics) based on those age-explicit estimates.

### 2 Constant survival and fecundity

We derive for each sex the constant survival probability from one age (or age class) to the next, such that the constant gives the same mean lifespan for the sex as its counterparts (i.e., those empirically observed age-dependent survival rates for the sex) would give. Under the assumption that each sex has a very large (ideally infinite) number of ages (or age classes), these constants can be theoretically derived; however, such assumptions are often violated in natural populations, and the use of such analytically derived constants may lead to notable deviations of the resulting lifespans from those calculated from data (the fewer the number of ages or age classes assumed for a sex, the more the analytically derived constant survival rate will underestimate the mean lifespan empirically derived for the sex). For biological realism, we first derive the constant survival probability for each sex analytically, and then keep *approximating* the constant computationally, until the difference between the empirically calculated mean lifespan for the sex and that the approximated constant yields becomes trivial (e.g., $10^{-4}$, see code availability for details). Below, we describe how these constant survival rates are theoretically derived as the basis of the approximations.

Let $l(a)$ be the probability that an individual of a given sex is alive at age (or age class) $a$, while $g(a)$ be the probability that the individual will survive from $a$ to $a+1$ (we assume ages or age classes are discrete). With empirically observed $l(a)$ and $g(a)$ for a sex, the mean lifespan $\bar{a}$ for the sex is

$$\bar{a} = \frac{\sum a \times l(a) \times [1-g(a)]}{\sum l(a) \times [1-g(a)]}$$

With $\bar{a}$ derived from data while assuming a large number of ages (or age classes) for the sex, we can analytically derive the constant survival probability $c$ from $a$ to $a+1$, such that $c$ will give the same mean lifespan for the sex as those age-dependent survival rates would give.

Let $g(a) = c$ (i.e., assuming age-independent survival rate for the sex), then by definition

$$l(a) = c^{a-1}$$

and thus the expected lifespan $a'$ for the sex is

$$a' = \frac{\sum a \times c^{a-1} \times (1-c)}{\sum c^{a-1} \times (1-c)} = \frac{\sum a \times c^{a-1}}{\sum c^{a-1}}$$

If a very large (ideally infinite) number of age classes for the sex is considered, then

$$a' = \lim_{a \to +\infty} \frac{\sum a \times c^{a-1}}{\sum c^{a-1}} = \frac{\frac{1}{1-c} + \frac{c}{(1-c)^2}}{\frac{1}{1-c}} = \frac{1}{1-c}$$

Now, let the empirically derived mean lifespan $\bar{a}$ be the same as the theoretically expected lifespan $a'$, and we have

$$\frac{1}{1-c} = \bar{a}$$

and thus we get

$$c = 1 - \frac{1}{\bar{a}}$$

as the constant survival rate for individuals of the sex. As these analytical solutions underestimate the empirically calculated mean lifespans for the sexes, we repeatedly increase these analytically derived constants for the sexes, until the approximations yield lifespans that are close enough to those empirically calculated for the sexes (see in the `R` code for details).

### 3 The hybrid version of the current model

Under the demographic processes assumed in the population, we can alternatively derive the patterns of kinship dynamics with a a Markovian-based hybrid approach: the patterns of kinship dynamics are derived from the analytically *expected* inter-individual relatedness under *simulated* changes of the age (class) compositions of a focal group over time, under the given sex- and age-(class-)specific survival probabilities of interest. Initially, the ages (or age classes) of individuals in each sex $\phi \in \{\alpha, \beta\}$ in the group are randomly drawn from the stable age (or age class) distribution for the sex (which is derived using the survival schedule of the sex). Let $l(K_\phi)$ be the probability that an individual of sex $\phi$ survives from age (or age class) 1 to $K_\phi$. As the survival schedules of both sexes depend only on individuals' ages, the stable age class distribution for the sex $\phi$ can be described by the probability $a(K_\phi)$ that an individual sexed $\phi$ is with age (or age class) $K_\phi$ at a given timestep, derived as

$$a(K_\phi) = \frac{l(K_\phi)}{\sum\limits_{K_\phi=1}^{C_\phi} l(K_\phi)} \tag{1}$$

With the stable age class distribution and the fecundity schedule for females, we can then derive the probability that in the focal group an individual sexed $\phi \in \{\alpha, \beta\}$ is born to a local female $i$ with age class $K_\alpha^i$ as

$$\begin{aligned} L_\phi(K_\alpha^i) &= \frac{(1-d_\phi)zF(K_\alpha^i)}{(1-d_\phi)z\sum\limits_{i=1}^{N_\alpha} F(K_\alpha^i) + d_\phi N_\alpha z \sum\limits_{K_\alpha=1}^{C_\alpha} a(K_\alpha)F(K_\alpha)} \\ &= \frac{(1-d_\phi)F(K_\alpha^i)}{(1-d_\phi)\sum\limits_{i=1}^{N_\alpha} F(K_\alpha^i) + d_\phi N_\alpha \sum\limits_{K_\alpha=1}^{C_\alpha} a(K_\alpha)F(K_\alpha)} \end{aligned} \tag{2}$$

where $z = \zeta$ when $\phi = \alpha$ (i.e., the individual is a female), while $z = 1 - \zeta$ when $\phi = \beta$ (i.e., the individual is a male). Therefore, the probability that this individual is native (or born) to the focal group is

$$L_\phi(\alpha) = \sum_{i=1}^{N_\alpha} L_\phi(K_\alpha^i) \tag{3}$$

With the stable age class distribution and the fecundity schedule for males, we can derive the probability that a male $i$ with age class $K_\beta^i$ claims his paternity in his group as

$$L(K_\beta^i) = \frac{\rho F(K_\beta^i)}{\rho\sum\limits_{i=1}^{N_\beta} F(K_\beta^i) + (1-\rho)N_\beta \sum\limits_{K_\beta=1}^{C_\beta} a(K_\beta)F(K_\beta)} \tag{4}$$

With (1), (2), (3), (4), we can then derive the probability that in the focal group a selectively neutral homologous gene will be copied from a female $i$ in age class $K_\alpha^i$, or a male $i$ in age class $K_\beta^i$, to an individual sexed $\phi \in \{\alpha, \beta\}$ native to the group as

$$H_\phi(K_\alpha^i) = \frac{1}{2} L_\phi(K_\alpha^i) \tag{5}$$

or

$$H_{\phi}(K^i_{\beta}) = \frac{1}{2}L_{\phi}(\alpha)L(K^i_{\beta}) \tag{6}$$

respectively.

We simulate inter-individual relatedness in the focal group over time with the probabilities of survival and inheritance. Let $\boldsymbol{R}_t$ be an $(N_\alpha + N_\beta) \times (N_\alpha + N_\beta)$ symmetric matrix capturing inter-individual relatedness at timestep $t$ in the focal group — initially we assume all the individuals in the group are not related to each other (but only to themselves), as denoted by the identity matrix $\boldsymbol{R}_0$. Then, the relatedness between individuals in the group at timestep $t+1$ (i.e., those survivors and/or newly-born individuals) can be derived with the mapping

$$\boldsymbol{R}_{t+1} = f(\boldsymbol{R}_t, \boldsymbol{Y}) \tag{7}$$

where $\boldsymbol{Y}$ is a sequence of survival and/or death events as the 'fates' of the females and males in the group at timestep $t$, which are simulated as a sequence of Bernoulli trials with the corresponding sex- and age-class-specific survival probabilities for these individuals. When mapping inter-individual relatedness at timestep $t+1$, we first classify each pair of individuals by their sexes and whether they are newly-established breeders or not given $\boldsymbol{Y}$ simulated. Then, we calculate the pairwise relatedness at $t+1$ based on $\boldsymbol{R}_t$ and the probabilities of inheritance derived in (5), and (6) — as we are interested in individuals' average relatedness to their groups as they age, we did not simulate the individually-explicit events of inheritance of the gene when doing the calculations, but calculated the analytically *expected* inter-individual relatedness instead.

More specifically, for each given $\boldsymbol{Y}$, we use $R_{t+1}(K^m_\phi > 1, K^n_\psi > 1)$, $R_{t+1}(K^m_\phi > 1, K^n_\psi = 1)$, or $R_{t+1}(K^m_\phi = 1, K^n_\psi = 1)$ to denote the expected relatedness between breeders $m$ and $n$ at timestep $t+1$ (i.e., the relatedness captured by the $m^{\text{th}}$ row and $n^{\text{th}}$ column in $\boldsymbol{R}_{t+1}$), where sexes are denoted by $\phi \in \{\alpha, \beta\}$ and $\psi \in \{\alpha, \beta\}$, and age class $K^m_\phi = 1$ (or $K^n_\psi = 1$) indicates that individual $m$ with sex $\phi$ (or $n$ with sex $\psi$) is newly-born in the group, while $K^m_\phi > 1$ (or $K^n_\psi > 1$) indicates $m$ (or $n$) is a survivor from the last timestep. Then, the relatedness between $m$ and $n$ at timestep $t+1$ is calculated as

$$R_{t+1}(K_\phi^m > 1, K_\psi^n > 1) = \boldsymbol{R}_t(m,n) \tag{8.1}$$

$$R_{t+1}(K_\phi^m = 1, K_\psi^n > 1) = \left[\sum_{g=1}^{N_\alpha} H_\phi(K_\alpha^g)\boldsymbol{R}_t(g,n) + \sum_{h=N_\alpha+1}^{N_\alpha+N_\beta} H_\phi(K_\beta^h)\boldsymbol{R}_t(h,n)\right] \tag{8.2}$$

$$\begin{aligned} R_{t+1}(K_\phi^m = 1, K_\psi^n = 1) = \Bigg[&\sum_{g=1}^{N_\alpha}\sum_{h=1}^{N_\alpha} H_\phi(K_\alpha^g)H_\psi(K_\alpha^h)\boldsymbol{R}_t(g,h) + \\ &\sum_{g=1}^{N_\alpha}\sum_{h=N_\alpha+1}^{N_\alpha+N_\beta} H_\phi(K_\alpha^g)H_\psi(K_\beta^h)\boldsymbol{R}_t(g,h) + \\ &\sum_{h=N_\alpha+1}^{N_\alpha+N_\beta}\sum_{g=1}^{N_\alpha} H_\phi(K_\beta^h)H_\psi(K_\alpha^g)\boldsymbol{R}_t(g,h) + \\ &\sum_{g=N_\alpha+1}^{N_\alpha+N_\beta}\sum_{h=N_\alpha+1}^{N_\alpha+N_\beta} H_\phi(K_\beta^g)H_\psi(K_\beta^h)\boldsymbol{R}_t(g,h)\Bigg] \end{aligned} \tag{8.3}$$

With initial $\boldsymbol{R}_0$, we can iterate (7) with (8.1), (8.2), and (8.3) for a given (large) number of timesteps $\mathcal{T}$, and derive the patterns of individuals' average sex- and age-class-specific relatedness to others in the focal group using the 'observed' inter-individual relatedness from the last $\tau$ timesteps ($\tau$ is also sufficiently large such that the number of observations for the last/highest age class can reliably capture the average relatedness the oldest individuals experience in the group).

### 4 Calculating relatedness of killer whales

We define females' local groups as their matrilines, sets of individuals with known lines of common maternal descent that are in near-constant social associations (Parsons et al., 2009). We use recently published genetic pedigrees (Kardos et al., 2023) to estimate individuals' relatedness to their matriline members within each census year. Relatedness is estimated from the pedigree using the *kinship2* R package (Sinnwell et al., 2014). We excluded individuals born prior to 1972 (due to uncertainty of their ages) and those without known first-order relatives.

### 5 Supplementary figures

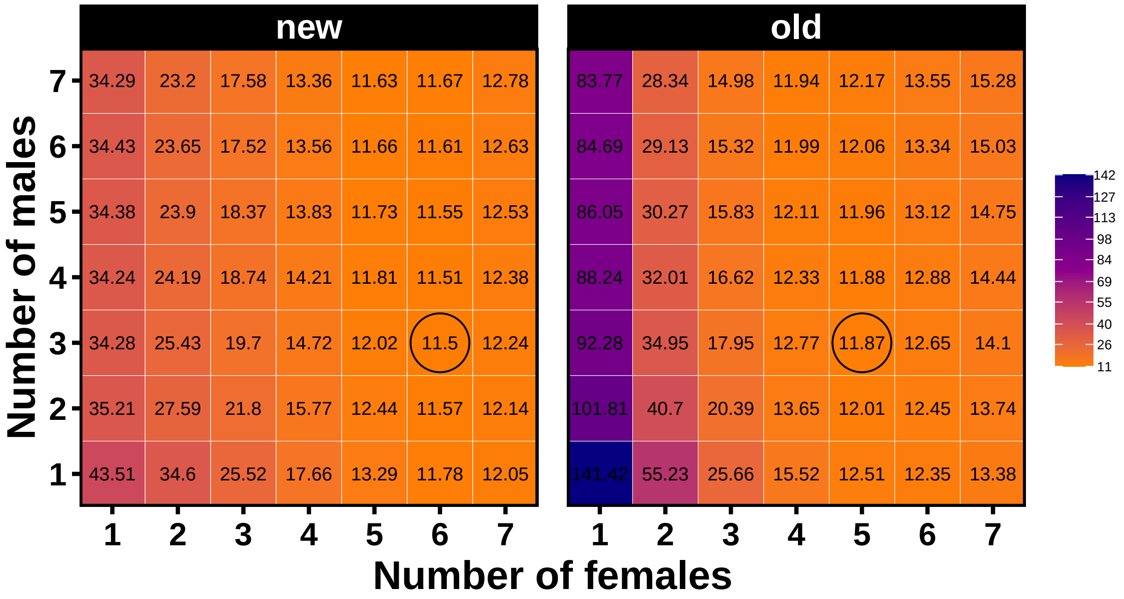


**Fig. A1** The sum of squared deviations calculated under different combinations of the numbers of females and males as inputs for both the new and old model of kinship dynamics. For each model separately, we calculated the predicted mean relatedness of males and females to their social group across the lifespan, for all possible combinations of the numbers of females and males taking integer values from 1 to 7, which define the smallest or largest group of 2 or 14 individuals, respectively, with sex ratio ranges from 1:7 to 7:1. The 'best-fit' numbers of females and males for the model in question are defined as those that minimise the sum of squared deviations between predicted relatedness values across both sexes and all age classes and those observed from individual whales at each age (marked by circles). In the killer whale example, the best-fit number of females is the same for both models (i.e., 3), while the best-fit number of males for the new and old models are 6 and 5, respectively.

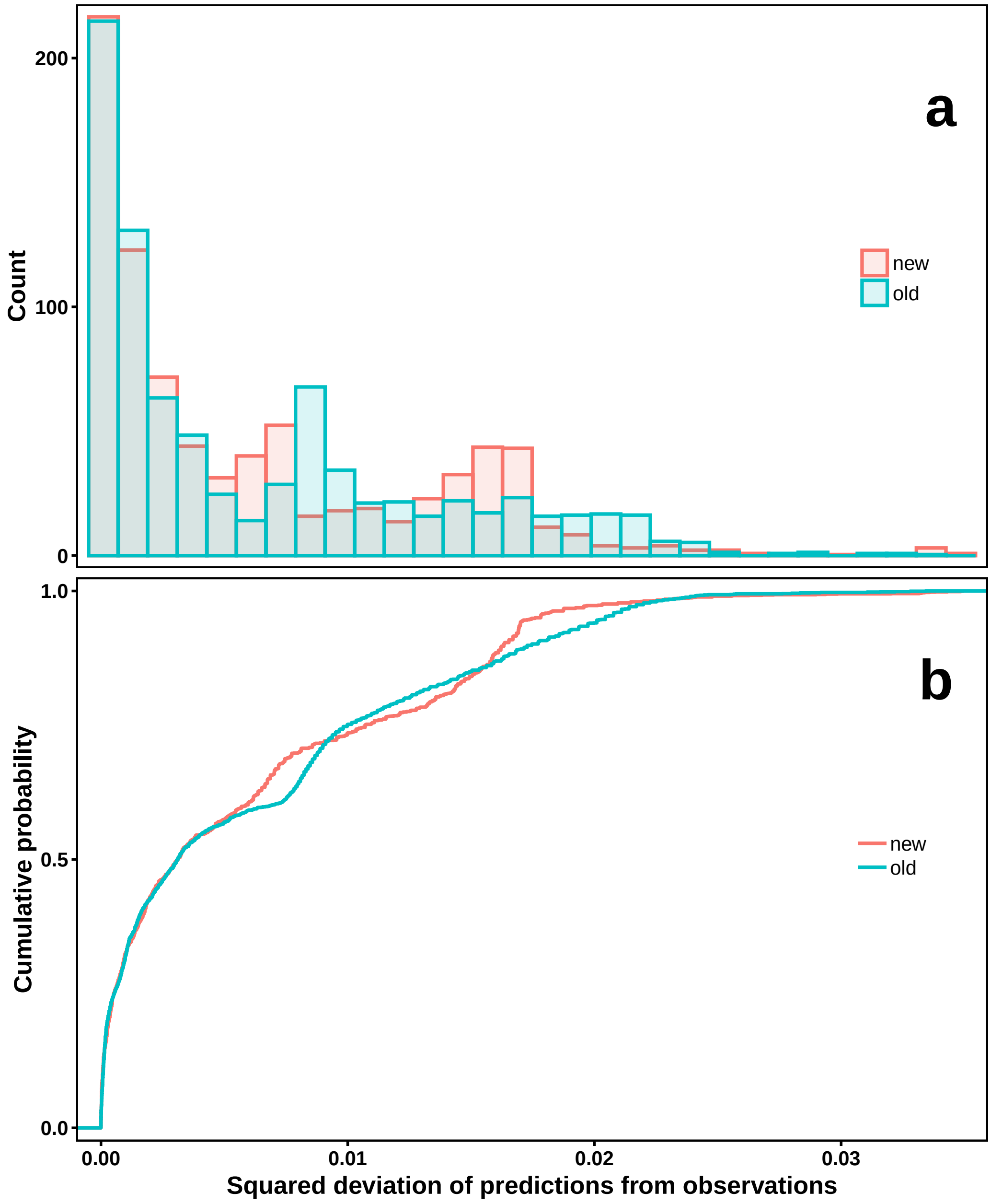


**Fig. A2** The distributions (a) and the empirical cumulative distributions (b) of the squared deviations of the sex- and age-specific predictions by the ‘new’ and ‘old’ model from empirical observations, with the ‘best-fit’ numbers of females and males for the model in question (Fig. A1).

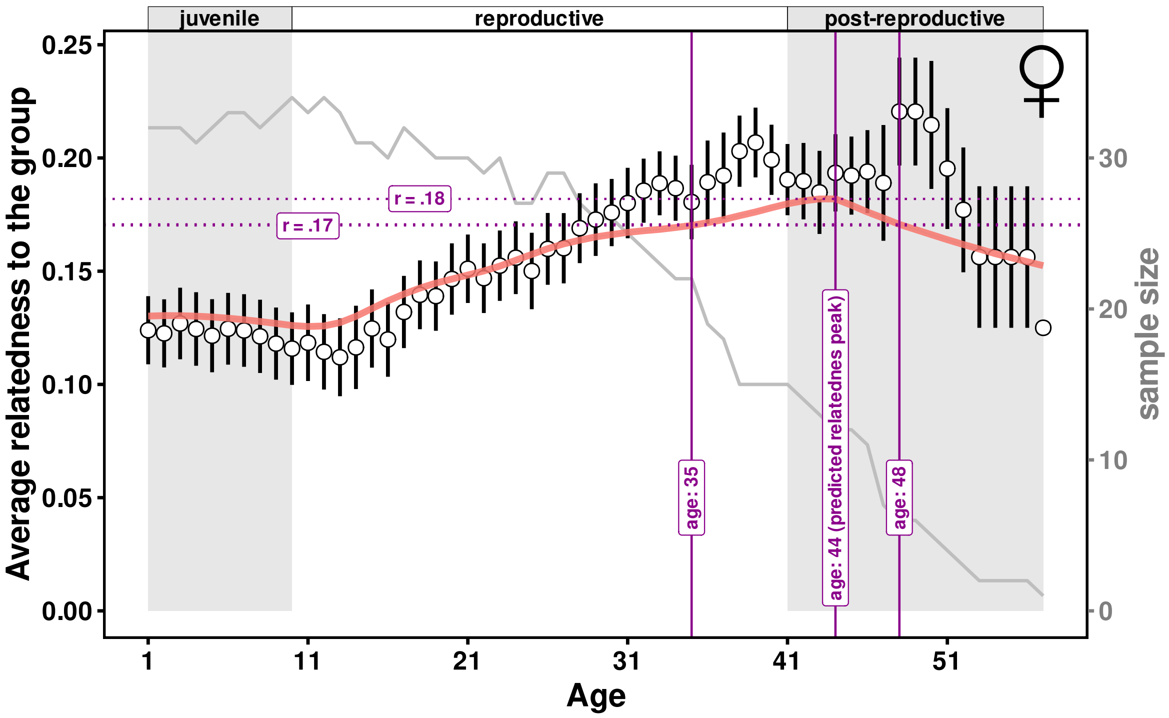


**Fig. A3** A (hypothetical) post-reproductive female dies at the age 48 (i.e., ca. 7 yeas after entering menopause) is predicted to be as strongly related ($r = .17$) to her group as she was when at the age of 35 — which is the (weighted) average age at death for females given their survival schedule, calculated as $\sum l(a) \times [1 - g(a)] \times a$ while assuming a stationary female age distribution [i.e., Fig. 1-d; here $l(a)$ is the probability that a female is alive at given age $a$, while $g(a)$ the probability that a female survives from $a$ to $a + 1$]. Here, the critical post-reproductive age 48 is derived by accounting for the expected number of years the female has been menopausal before her death, calculated as $\sum \frac{l(a)}{l(41)} \times [1 - g(a)] \times a$ (where $a > 41$, i.e., ages in post-reproductive lifespan), such that she is as strongly related to her group at her death at this critical post-reproductive age as she was when at age 35.
